# Rapid prediction of thermodynamically destabilizing tyrosine phosphorylations in cancers

**DOI:** 10.1101/2024.09.26.614998

**Authors:** Jaie Woodard, Zhengqing Liu, Atena Malemir Chegini, Jian Tian, Rupa Bhowmick, Subramanium Pennathur, Alireza Mashaghi, Jeffrey Brender, Sriram Chandrasekaran

## Abstract

Tyrosine phosphorylations are a prominent characteristic of numerous cancers, necessitating the use of computational tools to comprehensively analyze phosphoproteomes and identify potentially (dys)functional phosphorylations. Here we propose a machine learning-based method to predict the thermodynamic stability change resulting from tyrosine phosphorylation. Our approach, based on prediction of phosphomimetic delta-delta-G from structural features, strongly correlates with experimental mutational scanning cDNA proteolysis data (R = 0.71). We predicted the destabilizing effects of all 384,857 tyrosine residues from the Alphafold2 database. We then applied our approach to a pan-cancer phosphoproteomics dataset, comprising over 600 unique tyrosine phosphorylations across 11 cancer subtypes. We predict destabilizing phosphorylations in both oncogenes and tumor suppressors, where the former likely reflects a generalized relief of auto-inhibition or activating conformational change. We find that the number of circuit topological parallel relations with respect to residues contacting the phosphorylated site is greater for autoinhibited oncogenes than for other proteins (Wilcoxon p = 0.03). Utilizing an extreme gradient-boosting machine learning approach, we obtain an AUC of 0.85 for the prediction of autoinhibited phosphorylation states from circuit topological features. The top destabilized proteins from the pan-cancer data are enriched for chemical and oxidative stress pathways. Among metabolic proteins, highly destabilizing phosphorylations tend to occur in more peripheral proteins with lower network centrality measures (Wilcoxon p = 0.005). We predict 58% of recurrent tyrosine cancer phosphorylations to be destabilizing at the 1 kcal/mol threshold. Our approach can enable rapid screening of destabilizing phosphorylations and phosphomimetic mutations.

## Introduction

Protein tyrosine phosphorylation is frequently implicated in human cancers and other disease states (Peng et al. 2023; Reimand and Bader 2013) It is still a topic of some debate how much protein stability change contributes to alteration of protein function and pathogenesis. A common mechanism promoting disease is protein activation through relief of auto-inhibition or other conformational change, upon phosphorylation (Bayliss, Haq, and Yeoh 2015; Schlessinger 2003; Pufall and Graves 2002).

Deactivation, whereby alteration of interactions and/or alteration of protein stability could contribute to loss of function, may also promote disease, although such mechanisms are generally understudied (Bolduc et al. 2013; Geffen et al. 2021). Protein function can be lost due to destabilizing *mutations* (Woodard, Zhang, and Zhang 2021; Gao, Zhou, and Skolnick 2015; Cheng et al. 2023), begging the question of whether phosphorylation can be substantially destabilizing to the protein, intramolecularly. It has been proposed that phosphorylation can in fact commonly alter protein stability (Huang et al. 2019). Of phosphorylated residue types, tyrosine might be expected to have the broadest range of stability alteration, as hydrophobic interactions in the protein can potentially be affected through addition of a negative charge, leading to high destabilization of the protein. However, existing experimental studies lack substantial high-confidence information on stability change associated with tyrosine phosphorylation (Potel et al. 2021).

A plausible alternative to interrogating phosphorylation directly is to explore phosphomimetic modifications (Figure 1), whereby tyrosine, serine, or threonine is mutated to glutamate or aspartate, in a common experimental approach (Kliche et al. 2023; Sora et al. 2023; Luwang and Natesh 2018; Pearlman, Serber, and Ferrell 2011; Thorsness and Koshland 1987). Mutational scanning examples of phosphomimetic mutations in generic small proteins (not necessarily at phosphorylated sites) reveal that tyrosine to glutamate mutation tends to be much more destabilizing on average than mutation of either serine or threonine to glutamate (Tsuboyama et al. 2023). While computational modeling of phosphorylation and phosphomimetics has been attempted in the past, such studies have generally been limited to a small number of examples(Sora et al. 2023; Pérez-Mejías et al. 2020). One study found that phosphomimetic approaches can potentially be more accurate than direct phosphorylation modeling(Sora et al. 2023). Whole proteome scale experimental phosphorylation and phosphomimetic modeling has been attempted only for yeast (Bradley et al. 2024), likely due to questionable accuracy of such approaches. Determination of stability change due to phosphorylation at the proteome level presents experimental and computational challenges (Potel et al. 2021).

**Figure 1.**
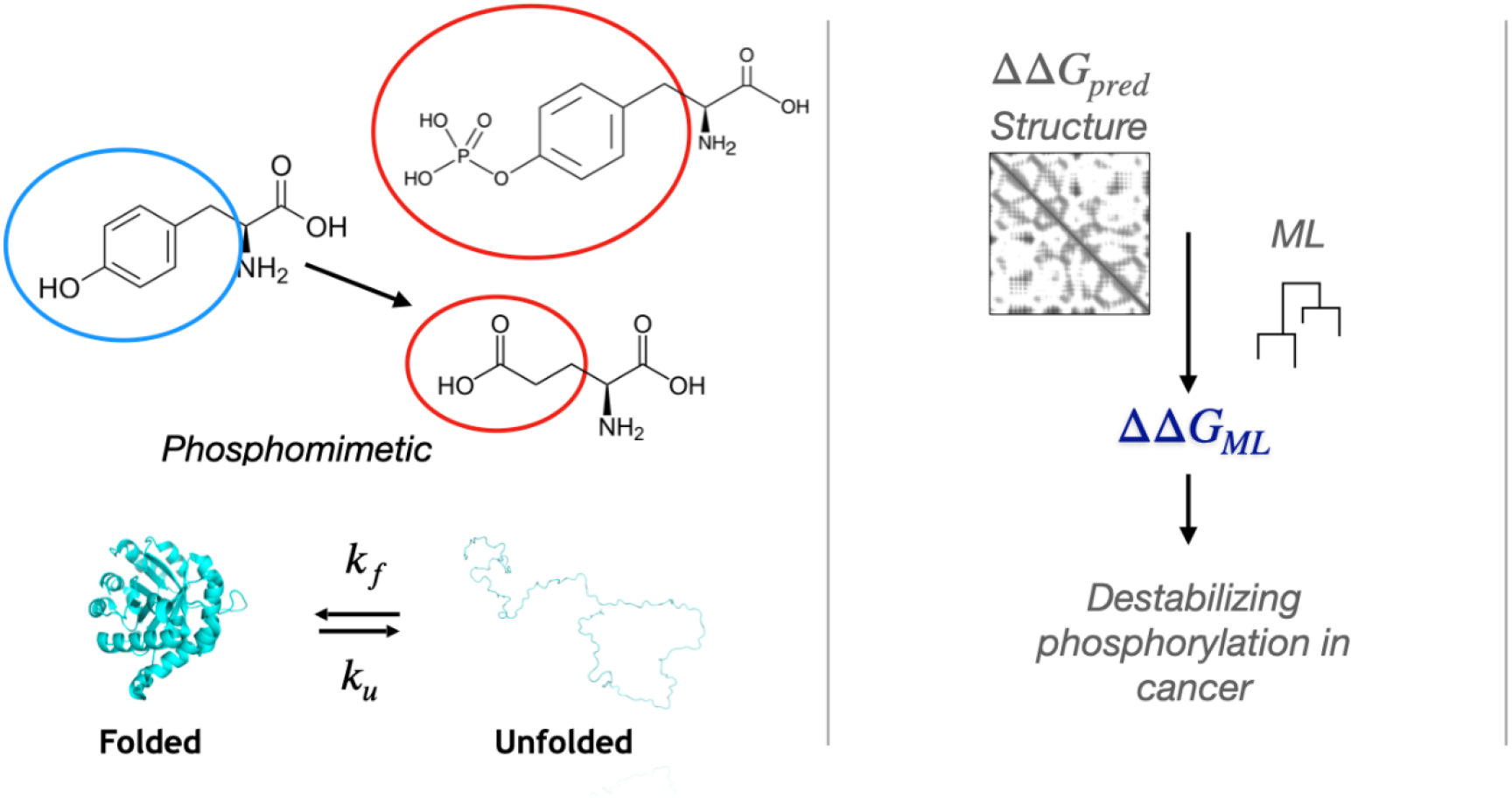
Outline of the approach presented in this work. Left: tyrosine is mutated to glutamic acid, to mimic the effect of phosphorylation. The folded vs. unfolded fraction of the protein in a two state model is estimated via computational approaches. Right: Structural features feed into a machine learning model to predict free energy change upon phosphorylation, ΔΔG. Our prediction method is applied to a pan cancer dataset to predict destabilizing phosphorylations in this context.

Protein thermodynamics is a useful framework for understanding how function can be lost due to destabilization. ΔG of folding determines the amount of protein in the folded vs. unfolded state at equilibrium in a two-state model(Tian et al. 2015; Fersht 1993). ΔΔG of mutation or phosphorylation is then the difference between ΔG of the modified protein and ΔG of WT, a measure of the change in protein stability. The sign convention is generally such that *positive* ΔΔG means that the amount of protein in the unfolded state at equilibrium increases, while the amount in the folded state decreases. While two-state models have limitations, they are generally considered a useful approach to quantifying changes in protein stability upon mutation or post translational modification.

The overarching goal of this study is to predict functional or maladaptive phosphorylation using ΔΔG prediction programs and machine learning (ML) (Figure 1). Previously, several groups have predicted functional phosphorylation using systems biology approaches, but they lack detailed structural information (Xiao et al. 2016; Reimand and Bader 2013; Smith et al. 2022; English and Torres 2022). Importantly, deep phosphoproteome experimental investigations have found that at least 75% of the proteome is phosphorylated, with tyrosine phosphorylations possessing distinct network properties from serine/threonine (Sharma et al. 2014). Databases are beginning to be developed which annotate structural features of phosphorylation and other PTMs within the protein structure (Ramasamy et al. 2020). Ultimately, our goal is to combine systems level approaches with structural approaches to develop a comprehensive picture of PTM signaling and static modification.

We present computational predictions of ΔΔG due to tyrosine phosphorylation as measured by three distinct methods: (1) phosphomimetics in FoldX and the methodologically similar program EvoEF, (2) ML based prediction from the Tsuboyama et al experimental dataset (Tsuboyama et al. 2023) and structural features, (3) Direct phosphorylation modeling in FoldX. We find high correlation between diverse methods of ΔΔG calculation. Using insights from these methods, we develop a ML method that rapidly predicts ΔΔG using a small set of easily obtainable features. We present several case studies, including proteins for which mutation to aspartate at the phosphorylated residue leads to clinically and experimentally evidenced disease. We focus in particular on the case studies of PI3K, for which destabilization of an inhibitory subunit by phosphorylation likely leads to activation, and HSP60, for which phosphorylation may alter the conformation of a nearby loop. We prioritize destabilizing tyrosine phosphorylations for further experimental study.

## Results

### Tyrosine phosphomimetic mutations are more destabilizing than serine or threonine

Our first goal was to determine whether certain types of phosphomimetic mutation were more destabilizing than others. Referencing mutational scanning experimental data from a proteolysis-based approach (Tsuboyama et al. 2023), we generated distributions of ΔΔG for mutation from serine, threonine, or tyrosine to glutamate. Such mutations are considered phosphomimetic, due to the addition of a negative charge mimicking phosphorylation. We found that tyrosine phosphomimetic mutations were more destabilizing on average, with a larger standard deviation than serine or threonine (Fig. 2). From visual inspection of corresponding structures, it seems likely that high numbers of contacts with the phosphorylated residue lead to greater destabilization. For instance, the most destabilized protein, the CD3 domain indicated by dark orange, contains tyrosine engulfed within the largest number of contacts with surrounding residues. This hypothesis that contacting residue number correlates with free energy change is explored in detail in the following sections.

**Figure 2.**
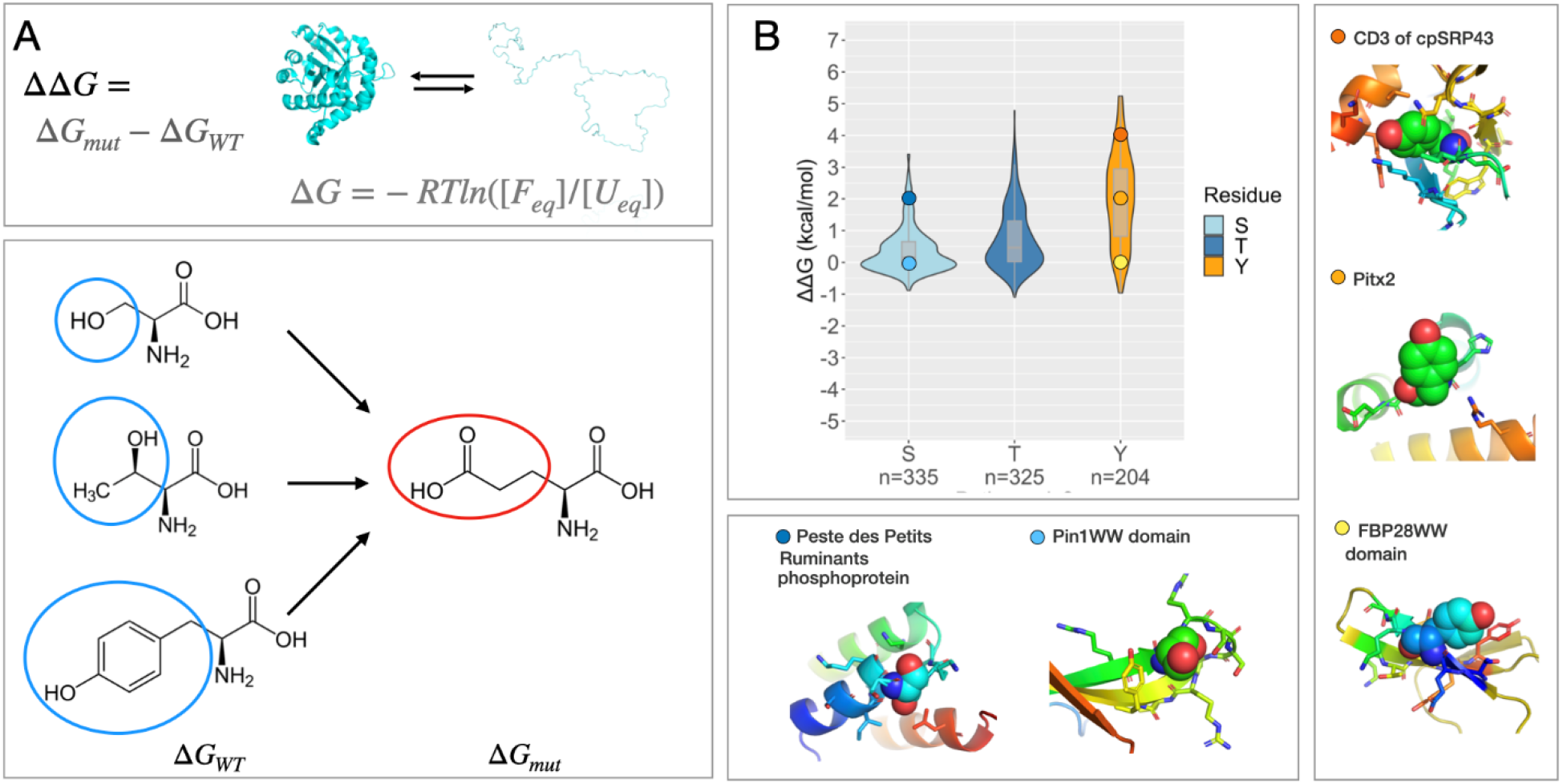
Determination of ΔΔG from Tsuboyama et al data. A) Diagram depicting phosphomimetic mutations references, along with equations for ΔG and ΔΔG. B) ΔΔG by residue type for mutation to glutamate from the Tsuboyama et al dataset.

### ΔΔG predictions for Tyrosine to Glutamate from structural features correlate with experiments

We wanted to see whether protein stability change calculations correlate with ΔΔG from Tsuboyama et al data. For calculations, we utilized two physical potential based methods with differing energy functions: FoldX (Schymkowitz et al. 2005) and EvoEF (Pearce et al. 2019). We calculated ΔΔG of phosphomimetic mutation from tyrosine to glutamate using the programs FoldX and EvoEF and found correlations of 0.56 and 0.53 with experimental values. Correlations for serine and threonine were more modest (0.36 and 0.30 for FoldX, and 0.17 and 0.24 for EvoEF, respectively). We also found a substantial correlation of experimental tyrosine to glutamate ΔΔG with number of residue contacts (r = 0.39). Strikingly, correlation between experimental ΔΔG and solvent accessible surface area was -0.65, suggesting that the amount of the residue accessible to the solvent is highly predictive of experimental stability change upon phosphomimetic mutation.

Correlation between FoldX and EvoEF values was 0.65. Figure S1 shows a scatterplot of FoldX vs. EvoEF ΔΔG with a color scale corresponding to experimental ΔΔG. FoldX overall tends to predict larger ΔΔG values than EvoEF, with the true value falling in between the two predictions. There appear to be two partially overlapping clusters in the data, with values below FoldX ΔΔG of about 2 kcal/mol having lower overall experimental energy and values above 2 kcal/mol having higher experimental energy. While FoldX ΔΔG is generally more predictive of experimental values than EvoEF, we note that the most destabilized point for EvoEF is also one of the most destabilized experimentally.

### Machine learning based on structural features improves tyrosine to glutamate ΔΔG correlation

We next sought to establish the predictability of phosphomimetic ΔΔG from structural features. We extracted several structural features in addition to EvoEF and FoldX predicted ΔΔG and used these as input for machine learning (Fig. 3A). The difference in ΔG between mutant and WT for natural sequences from Tsuboyama et al was considered as the predicted quantity. The features extracted were number of residues in contact with the phosphorylated residue (any atom within 5 Angstroms), solvent accessible surface area (SASA), and phi and psi angles (all calculated in PyMOL), and FoldX ΔΔG (Schymkowitz et al. 2005) and EvoEF ΔΔG (Pearce et al. 2019). Machine learning was carried out using Generalized Linear Modeling (GLM), Gradient Boosting Machine (GBM), and Extreme Gradient Boosting (XGBoost), using 5-fold cross validation. XGBoost was identified as the top performing model with cross validation correlation (R) of 0.71 (Fig. 3B). SASA was the top-ranking feature in feature importance utilizing this method, followed by FoldX ΔΔG and then EvoEF ΔΔG. Visual inspection reveals that phosphomimetic mutations that are predicted and confirmed to be highly destabilizing are in residues that are nested within a web of contacts with surrounding residues, while those that are least destabilizing tend to be in residues on the protein exterior, pointing outward, or in disordered regions (Fig. 3). Interestingly, both SO_1732 and bbc1 have a glutamate COO-group near the OH group of tyrosine, while the OH group of the less destabilized proteins is freely accessible to solvent (Fig 3C, 3D). Predictions for serine and threonine mutations to glutamate were also statistically significant with Tsuboyama et al data but lower correlation than for tyrosine. Full predictions are available in Dataset S1.

**Figure 3.**
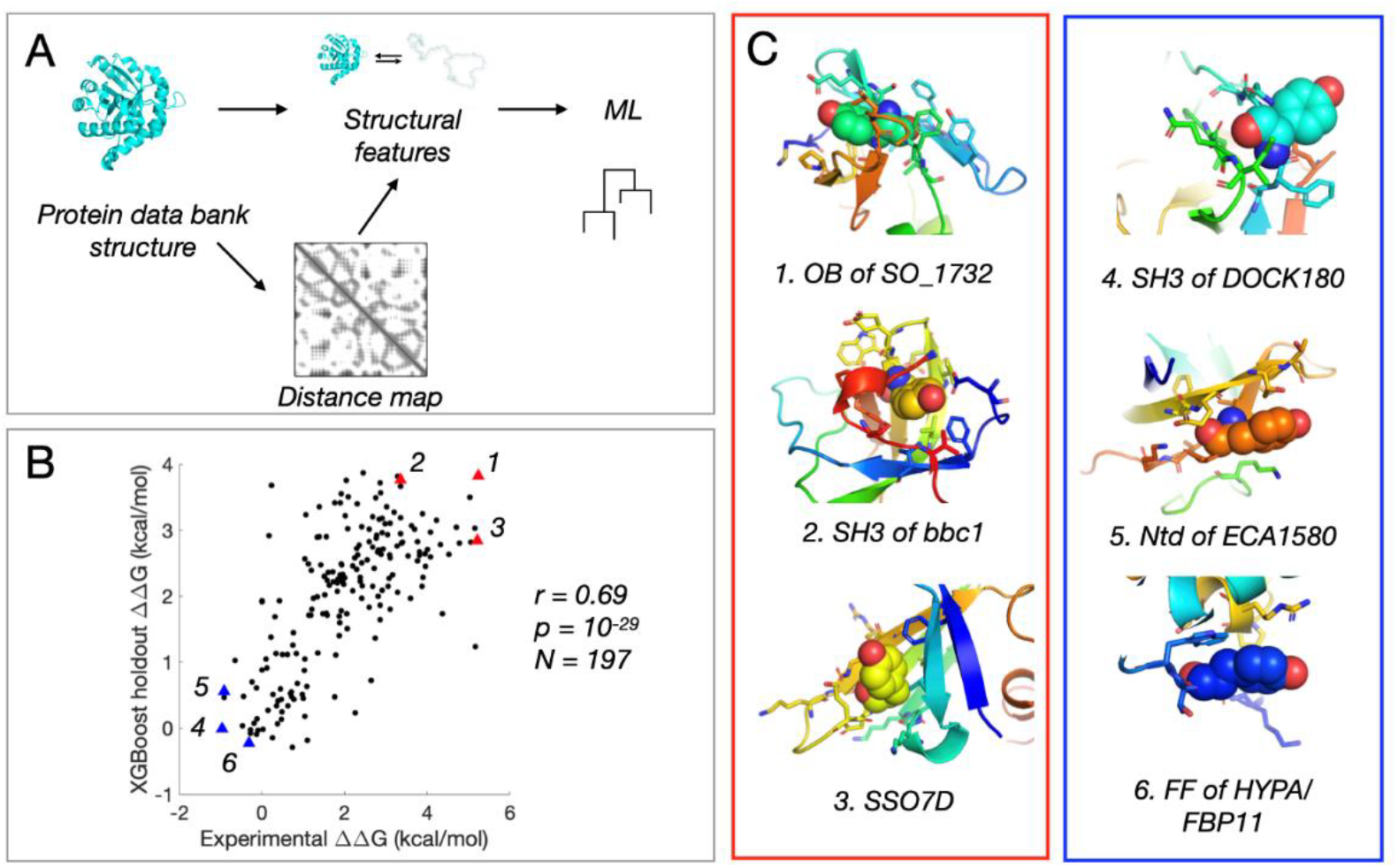
Predicting phosphomimetic stability (ΔΔG) based on structural features. A) Overview of the methods employed in this study. Protein data bank structures are used to extract protein structural features and contact maps, which are then input into machine learning algorithms. B) Scatter plot showing correlation (r = 0.71) between machine learned ΔΔG and experimental ΔΔG, for tyrosine phosphomimics. C) Representative structures from the boxed regions in (B), by box color. High ΔΔG residues are generally more interior and have more contacts.

### ΔTm values from Potel et al do not correlate with solvent accessible surface area

Surprisingly, we observed a general lack of consistency between solvent accessible surface area calculations and change in melting temperature upon phosphorylation, ΔTm, from Potel et al (Potel et al. 2021). Huang et al (Huang et al. 2019) recently developed a method to assess the thermodynamic stability change due to phosphorylation, the results of which claimed that a large fraction of proteins are significantly destabilized by physiological phosphorylations (S, T, and Y). Potel et al (Potel et al. 2021) built upon this method to develop an approach that they claimed was more rigorous, with which fewer phosphorylations were found to be thermodynamically destabilizing, and they highlighted some true positive case studies including an autoinhibition relieving phosphorylation of Fyn. We posited that if such results were to be meaningful, ΔTm would correlate substantially with SASA; phosphorylations further within the core of the protein would be more prone to destabilization. However, upon carrying out SASA calculations on residues from alphafold2 structures of the proteins, we found a correlation of less than 0.1 for each of serine, threonine, and tyrosine phosphorylation. We therefore reasoned that Potel et al data contains experimental and/or statistical complications that prevent it from being accurate enough to be physically reasonable and reliable. Therefore, we focused our efforts primarily on matching phosphomimetic data from the recent mutational scanning dataset by Tsuboyama et al (Tsuboyama et al. 2023).

### FoldX phosphorylation ΔΔG correlates with phosphomimetic approaches in a pan-cancer dataset

In order to compare phosphomimetic and direct phosphorylation methods, and as a useful application to cancer phosphorylation, we applied our methods to a pan-cancer dataset (Geffen et al. 2023). We found a quite high correlation of 0.80 between FoldX direct phosphorylation and XGBoost phosphomimetic method (Fig. 4). In the scatterplot, there are two clusters, separated by roughly XGBoost ΔΔG = 1 kcal/mol, an often-cited threshold for a destabilizing mutation to affect protein function. The boxplot in Fig. 4B shows that the 95^th^ percentile of phosphorylations below this threshold have lower ΔΔG of direct phosphorylation than the 25^th^ percentile of phosphorylations above this threshold. We note that it is possible that phosphomimetic results are a better representation computationally of phosphorylation than phosphorylation modeled in FoldX, as seen in a protein-protein interaction modifying phosphorylation in a recent computational study (Sora et al. 2023). Full data is available in Dataset S2. In prioritizing phosphorylations for future study, it is therefore informative to consider both scores. A complete table of correlations among EvoEF mutation, FoldX mutation, XGBoost, direct phosphorylation in FoldX, and SASA is available in the Table S1.

**Figure 4.**
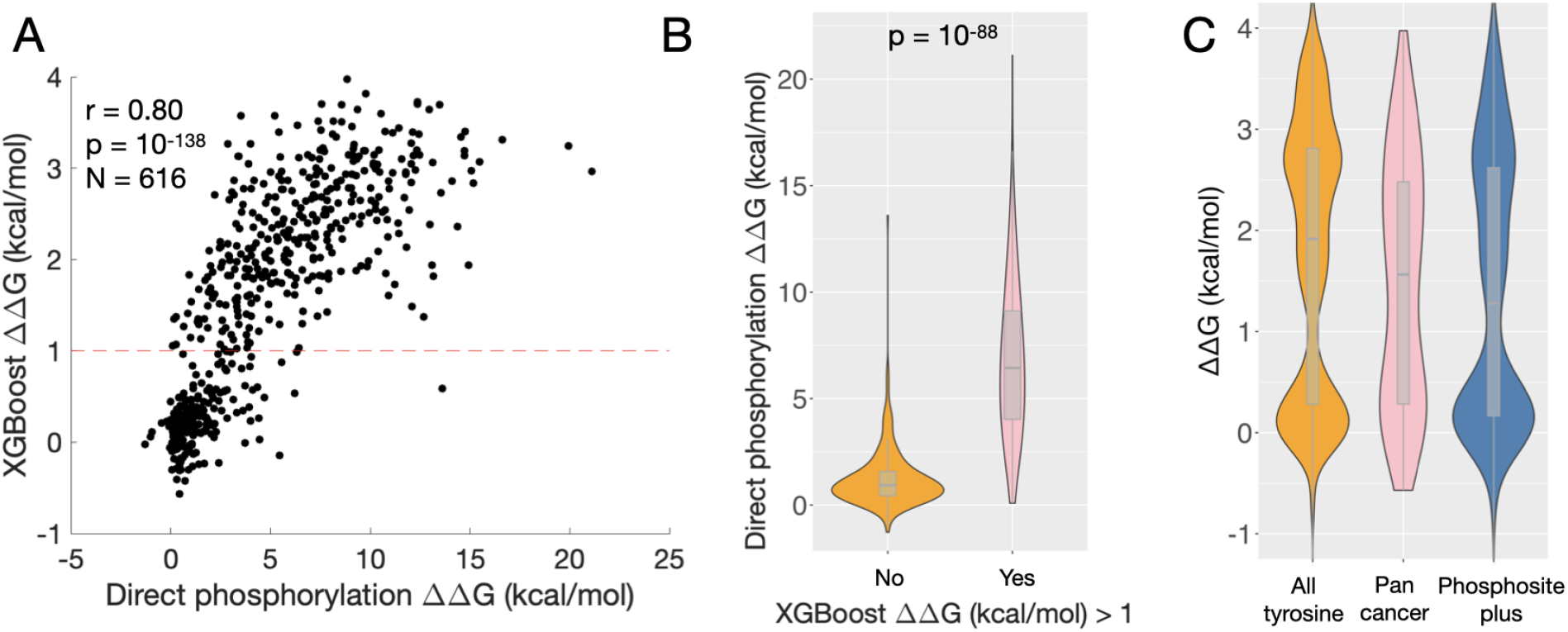
A rapid ML method to predict ΔΔG of phosphorylation. A) Relationship between XGBoost holdout predictions and direct phosphorylation ΔΔG in FoldX. B) Violin plots showing direct phosphorylation ΔΔG for XGBoost holdout ΔΔG less than or equal to, or greater than 1 kcal/mol. C) Violin plots showing distributions of rapid predicted ΔΔG for all Alphafold2 tyrosine residues, pan cancer data, and Phosphositeplus data.

### Feature reduction eliminating EvoEF and FoldX calculations improves speed without substantially compromising accuracy

We aimed to provide a rapid tool for predicting ΔΔG of phosphomimeitc change for tyrosine residues. To facilitate this, we decided to assess performance of the method upon eliminating the most computationally intensive steps: calculation of ΔΔG using EvoEF and FoldX, which demand about 10 and 24 hours of computational time to run, respectively, per 1000 mutations on large proteins such as those from the pan-cancer dataset. Upon eliminating these features, the calculation time reduced to approx. 3 minutes per 1000 mutations, and the performance only decreased to r = 0.67 (from r = 0.71 with EvoEF and FoldX). We therefore present our PyMOL feature/ML approach without energy change calculation as a rapid approach for estimating ΔΔG of phosphomimetic mutation. We make use of the rapid calculation method to predict ΔΔG for all 384,857 tyrosine residues from the Alphafold2 database. We find a distribution similar to that for tyrosine residues phosphorylated in cancers, with the distribution shifted slightly towards higher values for all residues vs. cancer (Fig. 4C, KS test p = 1×10^-9^). Additionally, we calculated rapid ΔΔG for the 36,468 human Phosphositeplus phosphorylated residues (Hornbeck et al. 2019), a database of all phosphorylations found in human, and found a distribution with a lower mean and median than for the cancer data (KS test p = 3×10^-4^), perhaps indicating a selection against disruptive and hard to access phosphorylations (Fig. 4C).

### Proteins destabilized by phosphorylation occupy peripheral network locations

To link protein structural properties with systems-level molecular network properties, we asked whether systems network centralities of proteins harboring highly destabilizing phosphorylation are different from those with less destabilizing phosphorylations. Specifically, we quantified centralities for metabolic reactions carried out by the proteins of interest, where reactions were counted as nodes and metabolites as edges. The network data was derived from a genome scale metabolic model, with centralities derived as part of a previous study(Smith et al. 2022). Metabolic networks were chosen as it is one of the most well studied and characterized cellular networks. We found that surprisingly degree, closeness, betweenness, and pagerank are all less for phosphorylations with ΔΔG greater than 1 kcal/mol vs. phosphorylations with lower ΔΔG (Figure S2). This suggests that cancer preferentially destabilizes proteins in network periphery, such as transporters, turning them off or destabilizing them. Perturbation of hub enzymes may be detrimental to the overall network function. Interestingly, XGBoost ΔΔG was not significantly different depending on gene essentiality identified using the Database of Essential Genes (DEG) (Luo et al. 2021; Zhang, Ou, and Zhang 2004), and modeled growth rate upon gene knockout was not significantly different for proteins with highly destabilizing phosphorylations (ΔΔG greater than 1 kcal/mol) vs. proteins with less destabilizing phosphorylations.

### Circuit topology distinguishes autoinhibition relieving phosphorylations in oncogenes

To further explore protein classes that are more destabilized in the pan-cancer data, we compared the stability effect of tyrosine phosphorylation on oncogenes and non-oncogenic proteins. Our methods predict that subsets of both tumor suppressors and oncogenes are highly destabilized by phosphorylation. For oncogenes, one mechanism by which destabilizing phosphorylation promotes tumorigenesis is through relief of autoinhibited states, activating the protein (Bayliss, Haq, and Yeoh 2015; Pufall and Graves 2002). We might ask whether it is possible to distinguish autoinhibited proteins from other proteins, and, more generally, oncogenes from tumor suppressors and other proteins. While oncogenes cannot be distinguished by ΔΔG alone (Fig. S4), we reasoned that this might be possible by considering the topology of the protein with relation to the phosphorylated residue. “Local” protein circuit topology (in contrast to “systems” network topology) provides a means to accomplish this. Given a contact, other contacts can be either in parallel, inverse parallel, series, or cross with this contact (Mashaghi, van Wijk, and Tans 2014). Additionally, local contact order, or the average distance along the chain to contacting residues, can be considered.

We find that the number of parallel relations is predictive of whether the protein is autoinhibited. Phosphorylations relieving autoinhibition in oncogenes are at residues exhibiting larger numbers of parallel relations relative to phosphorylated residues in tumor suppressors (Wilcoxon p = 0.03). We find high GBM machine learning-based predictability of autoinhibition relieving phosphorylations vs. phosphorylations in non-oncogenes (AUC = 0.85, MCC = 0.37). This suggests that autoinhibition-relieving phosphorylations share topological similarities, such as having many contacts on the “inside” of the autoinhibitory contact, such that relief retains a partially folded protein that is capable of function. Figure S5 illustrates parallel and inverse parallel relations for the simplified example of a hairpin and shows the result of removing contacts with many parallel relations, such that the protein core remains intact. Additionally, for the GBM classification of oncogene vs. non-oncogene, we achieve an AUC of 0.78 (MCC = 0.42), indicating that circuit topology is informative of protein oncogenicity after phosphorylation. Therefore, while free energy change itself does not distinguish oncogenes from other proteins, topology is a good predictor of this, facilitating a more complete picture of how phosphorylation leads to pathogenesis in cancer.

### Functional enrichment of destabilized genes

To assess the preference of protein destabilization to specific protein functional categories, we determined functional enrichment for the top 20% of genes destabilized by tyrosine phosphorylation in comparison to all other proteins (Figure S6). We found multiple enriched categories related to the epithelial-to-mesenchymal transition, known to be important in cancer-related processes, including metastasis, immune evasion, cell-cell adhesion, and wound healing. In addition, we found multiple categories related to cellular oxidative stress, including glutathione metabolic process and cellular response to chemical stress. Pyruvate metabolic pathway was also enriched, which is related to the well-known Warburg Effect in cancers. Overall, this functional enrichment analysis highlights several pathways known to be highly important in cancer and demonstrates that destabilizing phosphorylation likely influences cancer progression.

### Case studies of destabilizing phosphorylations

As a case study, we examined phosphorylation within the p85 subunit of Phosphatidylinositol 3-kinase (PI3K), a critical regulator of cell growth and survival. Of proteins with the most destabilizing tyrosine phosphorylation (in the top 90^th^ percentile), the PI3K inhibitory p85α subunit (PIK3R1 gene) stands out in that the relative orientation of two phosphorylated tyrosine residues closely resembles that of the receptor tyrosine kinase (Fig. 5A-B). Mutations within the PI3K domain that is phosphorylated are known to cause cancer. Phosphorylation of Y580 (orange in Figure 5A), in addition to being implicated in cancer, occurs in normal cells via the insulin receptor (Hayashi et al. 1993). It has been proposed that Y580 is a passenger phosphorylation in cancer (Nussinov et al. 2021). We suggest the alternative hypothesis that phosphorylation of this residue relieves inhibition of the interacting catalytic subunit, activating PI3K. Such activation may be functional in normal cells but is likely co-opted in cancer cells to promote cell growth and proliferation. We note that our phosphomimetic methods predict that Y580 phosphorylation is at the 91^st^ percentile of destabilizing phosphorylations overall (Table 1). Joint phosphorylation of the nearby tyrosine 452 in cancer may increase this effect.

**Table 1.** Phosphorylation case studies.

| Gene | Phosphorylated residue | Similar mutation(s) | Mutation phenotype | GBM $\Delta\Delta G$ percentile | Protein-protein interactions |
| --- | --- | --- | --- | --- | --- |
| PIK3R1 | Y580, Y452 | Mutations in same domain | Cancer, insulin sensitivity (Tsay and Wang 2023) | 91, 42 | Domain interacts with catalytic subunit (Nussinov et al. 2021) |
| PTPN11 | Y62 | Y62D | Noonan syndrome (increased cancer risk) (Martinelli et al. 2012) | 83 | NA |
| HBB | Y131 | Y131D | Anemia (Lorkin et al. 1974) | 94 | NA |
| KRT14 | Y129 | Y129D | Dowling-Meara Epidermolysis Bullosa Simplex (Chan et al. 1996) | 27 | Coiled coil interaction |

**Figure 5.**
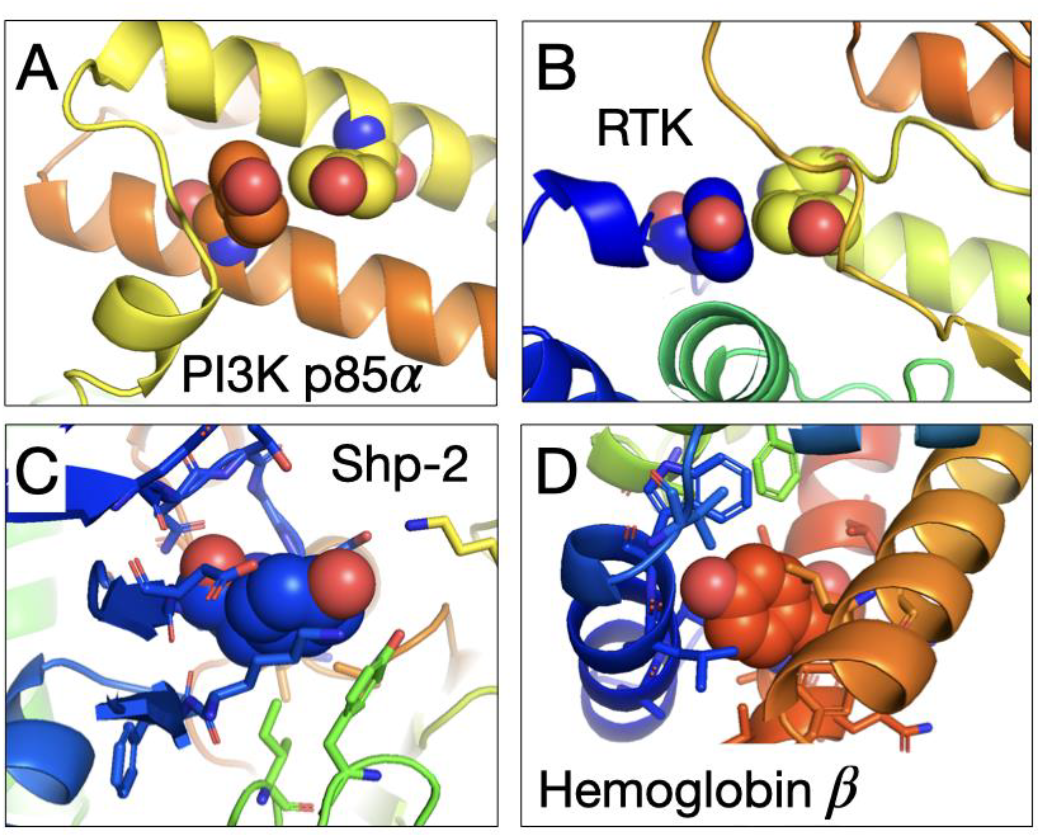
Protein destabilizing phosphorylation case studies. A) PI3K. B) RTK. C) PTPN11. D) HBB.

A second case study was cancer-related phosphorylation of the protein HSP60. HSPD1 or HSP60 is a member of the heat shock protein family. Hsp60 can be phosphorylated at Y227 and Y243, which is crucial for its surface activation. We find HSPD1 Y243 as one of the top hits using our XGBoost model, in the 93rd precentile. Phosphorylation occurs at a residue in contact with a large loop, which may exhibit phosphorylation-dependent conformational changes. HSP60 is present in the plasma membrane of human leukemic CD4+ T cells and is phosphorylated by protein kinase A (PKA). Hsp60 physically associates with Histone 2B (H2B) in the dephosphorylated form. Protein kinase A-catalyzed phosphorylation of HSP60 regulates its attachment to H2B (Khan et al. 1998). By contrast, PKA-catalyzed phosphorylation causes dissociation of H2B from HSP60, potentially due to its destabilizing effect.

We next wanted to see how evolutionary mutations at cancer-phosphorylated sites affect protein function. While we utilized mutation to glutamate as a phosphomimetic in this study, in evolution, tyrosine is more frequently mutated to aspartate, given similarity in codon usage rules. We mined the Humsavar database of rare variants for cases of mutation of tyrosine residues phosphorylated in cancer to aspartate. We found three such cases (Table 1, Figure 5C-D). All three are associated with pathogenicity to different extents. Of the three, two mutations are at sites of high predicted destabilization due to phosphorylation (in the top 20%); the third likely affects protein-protein interactions. Notably, for the proteins hemoglobin beta (HBB) and Shp-2 (PTPN11) (Figure 4C), the Tyr to Asp mutation confers increased cancer risk in genetic disease (Martinelli et al. 2012). Histograms of ΔΔG values with locations of example proteins labeled are shown in Figure S3.

## Discussion

Phosphomimetic approaches are commonly employed experimentally to interrogate functional phosphorylations. Numerous studies have demonstrated that phosphomimetics are often suitable representations (Kliche et al. 2023; Luwang and Natesh 2018; Pérez-Mejías et al. 2020) although there is still some debate (Pérez-Mejías et al. 2020; Jonson and Petersen 2001). This suggests that phosphomimetics may also be employed computationally, taking advantage of optimized strategies to model mutation between amino acid types (Sora et al. 2023).

With our simple machine learning method incorporating ΔΔG predictions from two highly employed energy function-based programs, we obtain performance on phosphomimetic mutations approaching the state of the art for the best GNN approaches ((Dieckhaus et al. 2024), r = 0.76 for the Tsuboyama dataset). Our method employed an XGBoost model with readily obtainable and mechanistic features, in addition to calculations that represent composites of mineable energy terms, and so is easy to interpret. SASA ranked as the top feature for tyrosine to glutamate mutation. We find high correlation between the XGBoost phosphomimetic based approach and direct phosphorylation modeling in FoldX, suggesting that both may be reasonably accurate representations of ΔΔG of phosphorylation. We suggest considering both predictions when prioritizing phosphorylations for further study. We also find that the phosphorylations predicted by phosphomimetics to be most destabilizing can be rationalized by examination of the protein structure. In addition to simply modeling the phosphomimetic mutation with traditional ΔΔG prediction tools, we employed machine learning strategies to recapitulate experimental results of a widely used mutational scanning dataset. Previous experimental studies did not identify a difference in ΔΔG of tyrosine phosphorylation vs. serine or threonine. However, such studies did not include a large number of high confidence tyrosine phosphorylations, and results were likely skewed towards less destabilizing phosphorylations. Experimental ΔTm of phosphorylation from the Potel et al dataset (Potel et al. 2021) did not correlate with SASA and ML model predictions. It is possible that experimental complications still exist in such data, although we cannot rule out that phosphomimetics may not be accurate representations of phosphorylated states generally. It is also possible that larger proteins possess properties that preclude forecast based on the smaller proteins of the Tsuboyama et al dataset. However, the identification of visually “obvious” destabilizations by our phosphomimetic approach suggests that this approach may be an adequate strategy.

We predicted that many phosphorylations in human cancers destabilize the protein beyond the often-cited threshold of 1 kcal/mol, beyond which mutations often affect protein function. While most proteins have stabilities of a few kcal or more, kinetic factors and the possibility of local unfolding push the threshold to lower values. Our results suggest that tumor suppressors can be depleted by thermodynamically destabilizing phosphorylation. Drugs that prevent phosphorylation or enzyme replacement therapy with a stable protein that cannot be phosphorylated may be beneficial in such cases. Among the cancer associated phosphorylations, the p85 subunit of PI3K illustrates a case in which the structural motif found in receptor tyrosine kinase (RTK) is reused in a different context. Similar to the relief of auto inhibition within the same protein chain via partial unfolding seen in RTK, destabilization of p85 likely abolishes the chain’s role in inhibition of the catalytic subunit. In both cases, two interacting tyrosine residues with similar respective orientations are both phosphorylated. However, the structural details surrounding the tyrosine-tyrosine interaction are quite different in these two cases. This is one example of a structural motif that is transferrable to other contexts, a common occurrence in structural biology.

While this study focused on applications to cancer, we note that destabilizing phosphorylation will likely be applicable to other disease states, including diabetes, as forecast by the PI3K case. However, there may be selection against destabilizing phosphorylations in healthy individuals (Bradley et al. 2024),also evidenced by the relative lack of destabilization in Phosphositeplus data (Fig. 4C). Hence, cancers may be particularly prone to harbor destabilizing phosphorylations. Further, while we focused on single phosphorylations, a limitation of our study is that many proteins contain multiple phosphorylation sites that may interact to determine overall effects. Autoinhibited proteins, for instance, often have two tyrosine phosphorylations side by side, as we also saw for PI3K. Phosphorylation at both of these sites likely leads to enhanced local or global destabilization of the domain. It should be possible to model such multiple phosphorylations using either phosphomimetics or direct phosphorylation approaches in the future. An important question moving forward is whether other post translational modification types may also exhibit stability alteration. A future extension would be to acetylation, for which mimetics also exist. In fact, stability alteration due to acetylation has been proposed and verified experimentally (Narita, Weinert, and Choudhary 2019; Incani et al. 2014; Shimizu et al. 2021). Methylation may be an example for which prediction is less straightforward, but results may also be determinable by rigorous computational approaches. Finally, a thermodynamic and structural analysis of PTMs such as glycosylation and SUMOylation via molecular dynamics and topological methods (e.g., (Woodard, Iqbal, and Mashaghi 2022) would likely reveal interesting results.

In sum, based on our extensive ML analysis, we find that ΔΔG of tyrosine phosphomimetic mutations can be accurately predicted. We identify protein circuit topology as a tool to discover autoinhibited states and oncogenicity. We propose adding destabilization of tumor suppressors due to phosphorylation to the repertoire of effects with potential to affect cancer progression. We prioritize predicted destabilizing phosphorylations for further experimental study.

## Methods

### Machine learning and ΔΔG of phosphomimetic mutation

Machine learning was carried out in order to predict ΔΔG of phosphomimetic mutation from structural features and energy calculations. FoldX was carried out with pH = 7.3 and ion strength = 0.15, with — pdbHydrogens = true and —water = IGNORE. PDB structures were aligned to the experimental protein sequence using the Needleman-Wunsch algorithm. In preliminary analysis, we found that addition of PSSM sequence conservation data to the machine learning model did not improve model performance, although performance of PSSM alone was near r = 0.4. Additionally, we considered the performance of models incorporating only structural features not including EvoEF and FoldX calculations, to achieve a speed-up of calculations with minimal decrease in accuracy. Machine learning was carried out in Python using the scikit learn package and 5-fold cross validation, with a grid search to identify top hyperparameters, where hyperparameters leading to a gap in cross validation predictions were avoided. Hyperparameter optimization is shown in Tables S3 and S4. The best hyperparameters for both the rapid method and the regular method were: {‘n_estimators’: 150, ‘learning_rate’: 0.01, ‘max_depth’: 3}.

### Application to the pan-cancer dataset

In order to utilize our approach in the context of cancer phosphorylation, we applied our machine learning predictions to a recent pan-cancer dataset ((Geffen et al. 2023). Sequences flanking the phosphorylation site were aligned to fasta sequences using the <u>Needleman Wunsch</u>algorithm, with gap and extension parameters altered to promote proper alignment; we found that 21% of aligned residue numbers did not conform with the numbering system employed in the publication. Phosphorylations were considered individually in cases where multiple phosphorylations occurred in the same protein. Alphafold2 (Jumper et al. 2021) structures were referenced. Features were extracted as above. In addition, direct phosphorylation modeling, or phosphorylation performed with the FoldX energy function, was carried out in FoldX (Schymkowitz et al. 2005), and correlation between the two methods of ΔΔG determination was assessed.

### Metabolic gene centralities

We sought to determine whether certain types of network locations were prone to harbor destabilizing phosphorylation. Degree, betweenness, closeness, and pagerank of metabolic reactions were extracted from (Smith et al. 2022), based on the Recon1 Genome Scale Metabolic Model (GEM) (Rolfsson, Palsson, and Thiele 2011). For genes that catalyzed multiple reactions, centralities were averaged over reactions for the gene of interest. Boxplots of centrality were generated for proteins predicted to be destabilized or not by greater than 1 kcal/mol.

### Circuit topology of autoinhibited proteins and non-oncogenes

We aimed to determine if it was possible to distinguish autoinhibited oncogenic proteins from non-oncogenes on the basis of topological parameters from circuit topology. 26 proteins for which tyrosine phosphorylation relieves autoinhibition were identified via a literature search. In cases where a protein contains more than one such phosphorylation, the phosphorylation with the greatest amount of literature evidence and frequency was chosen for analysis. All 26 proteins showed some literature evidence of oncogenicity. Local circuit topology and contact order were extracted for phosphorylated residues, as described previously (Woodard, Iqbal, and Mashaghi 2022; Moes et al. 2022), for the 26 proteins and proteins without CancerMine oncogene annotation from the pan-cancer dataset (“non-oncogenes”).

Briefly, circuit topology descriptions consist of parallel, inverse parallel (see Figure S5), series, and cross relations, where local circuit topology counts the number of each type of relation relative to any contact that is formed with the residue of interest. Distributions were compared, and GBM machine learning was carried out in Python using scikit learn. A grid search was performed to find optimal parameters, using 5-fold cross validation. For autoinhibited proteins, these were: {‘learning_rate’: 0.1, ‘max_depth’: 3, ‘n_estimators’: 100}, and for oncogenes vs. non-oncogenes, they were: {‘learning_rate’: 0.2, ‘max_depth’: 7, ‘n_estimators’: 100}. Following the grid search, an independent 5-fold cross validation was carried out to obtain final performance metrics. Hyperparameter optimization is shown in Tables S5-S6.

## Data Availability

Code is available on github: https://github.com/jaiewd/phosphorylation

## Acknowledgements

We thank Dr. Sumaiya Iqbal, Dr. Matthew O’Meara and members of the Chandrasekaran lab for helpful discussions. This work was supported by faculty start-up funds from the University of Michigan (UM), Camille and Henry Dreyfus Foundation, the Rogel Cancer Center at UM, R35 GM13779501 from NIH to SC. JW is supported by the NIH Predoctoral and Postdoctoral Multidisciplinary Training Program in Benign Kidney, Urology, and Hematology Disease, University of Michigan (U2C/TL1 Training grant U2CDK129445/TL1DK136046).

---

**Figure S1.**
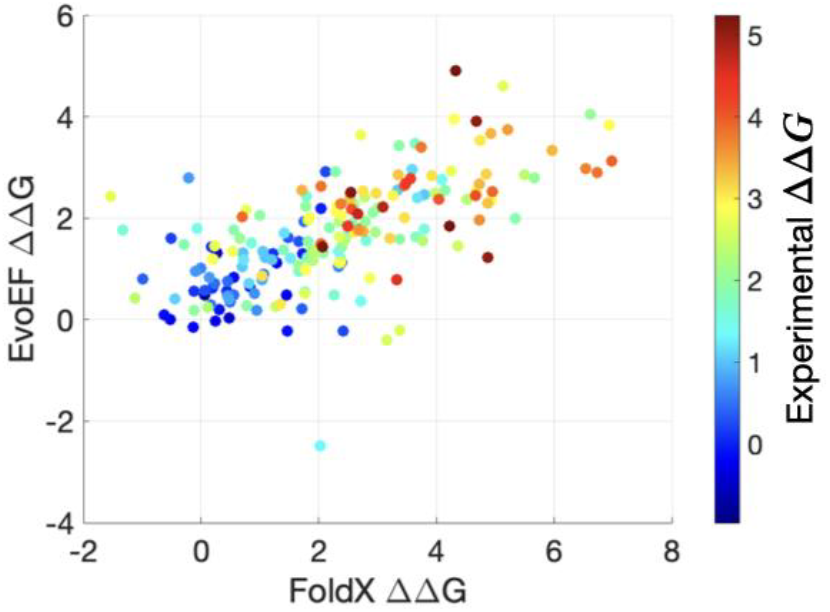
Predicted and experimental energy in units of kcal/mol, for Tsuboyama et al dataset for tyrosine to glutamate mutation. EvoEF ΔΔG vs. FoldX ΔΔG is plotted, with the color scale corresponding to experimental ΔΔG.

**Figure S2.**
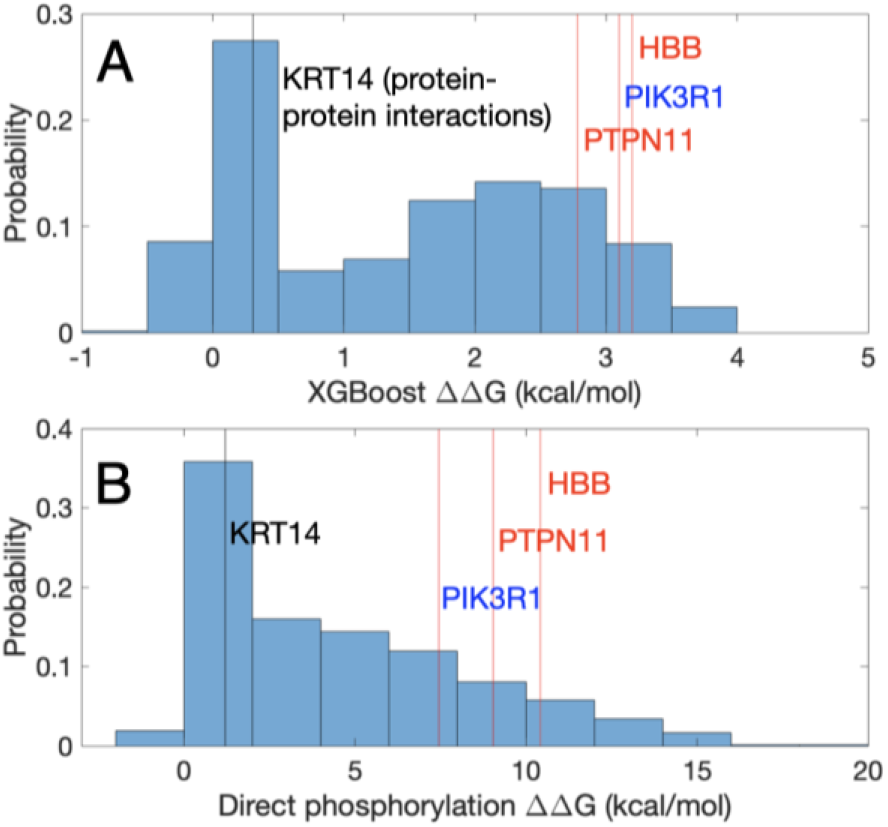
Metabolic network (GEM) centralities for machine learning predicted ΔΔG less than or equal to, or greater than, 1 kcal/mol. P value corresponds to a Wilcoxon test. Proteins predicted to be highly destabilized by phosphorylation consistently have lower network centralities.

**Figure S3.**
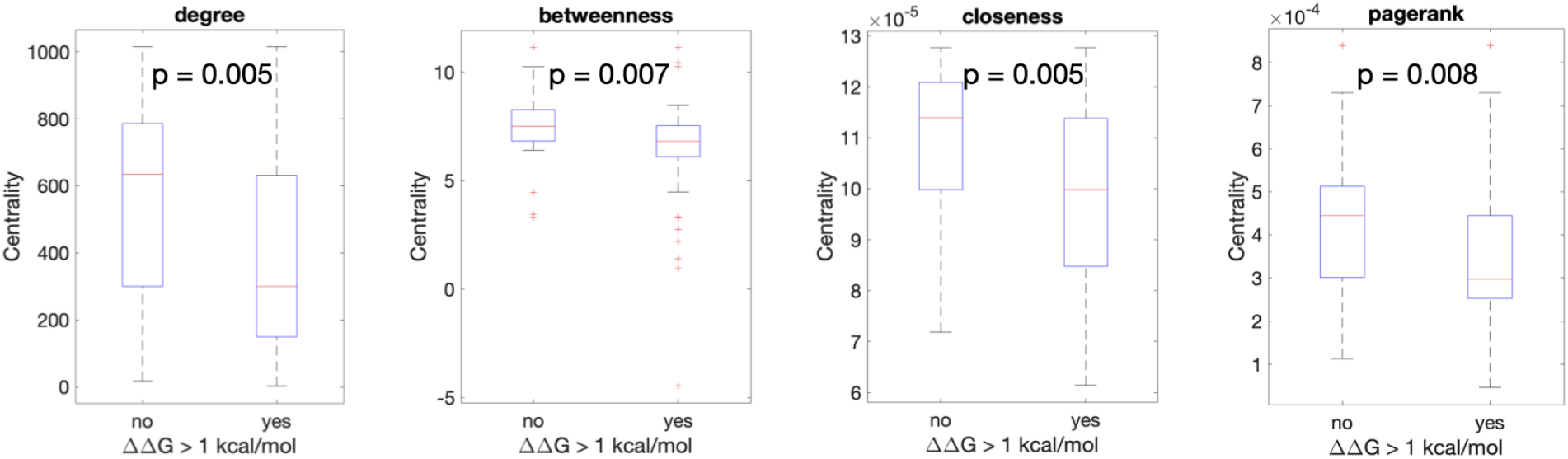
Histograms of stability change, for XGBoost predictions (A) and direct phosphorylation in FoldX (B). Genes from the table in the main text are indicated.

**Figure S4.**
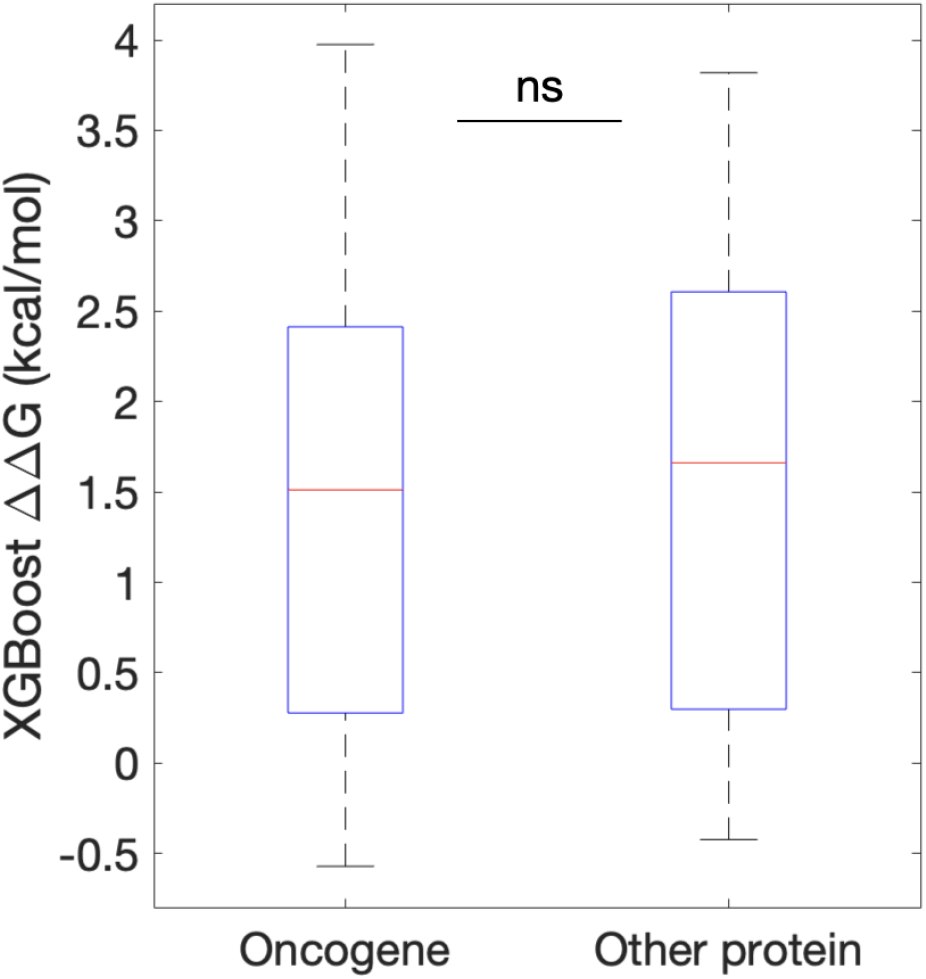
Predicted free energy change distribution for phosphorylations belonging to oncogenes and belonging to non-oncogenic proteins. Difference is non-significant by t-test or Wilcoxon test.

**Figure S5.**
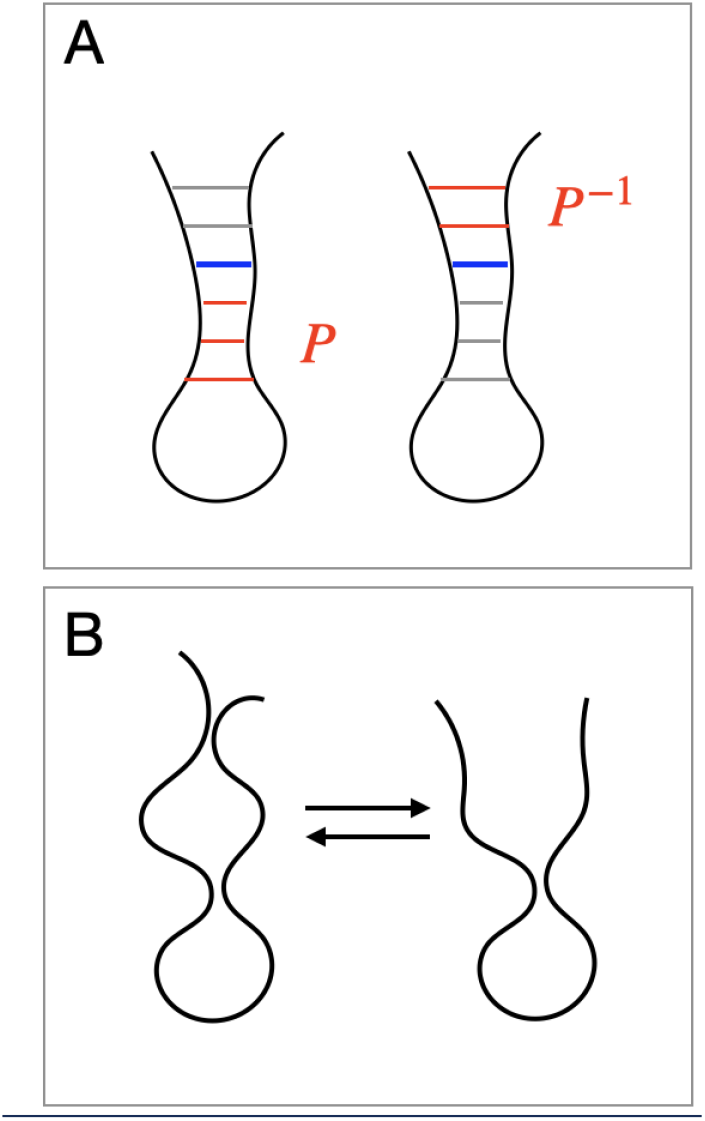
Circuit topology relations. A) Parallel and inverse parallel relations for the simplified example of a hairpin. B) Result of deleting contacts with many parallel relations.

**Figure S6.**
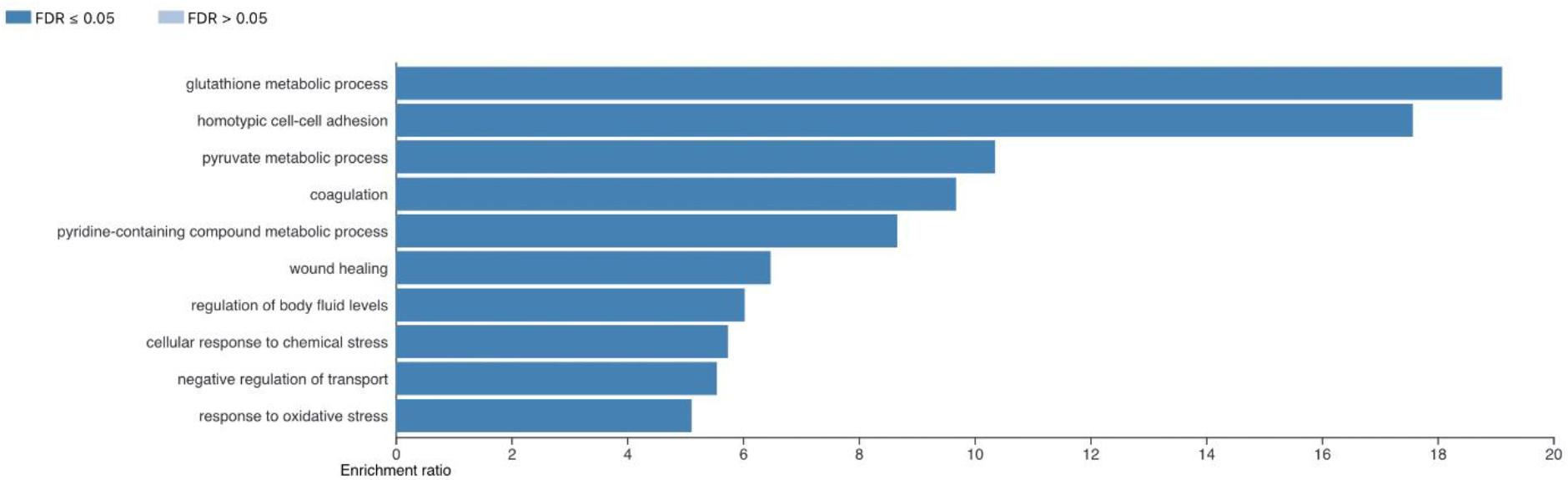
Top functional enrichment categories, calculated in GSEA.

**Table S1.**
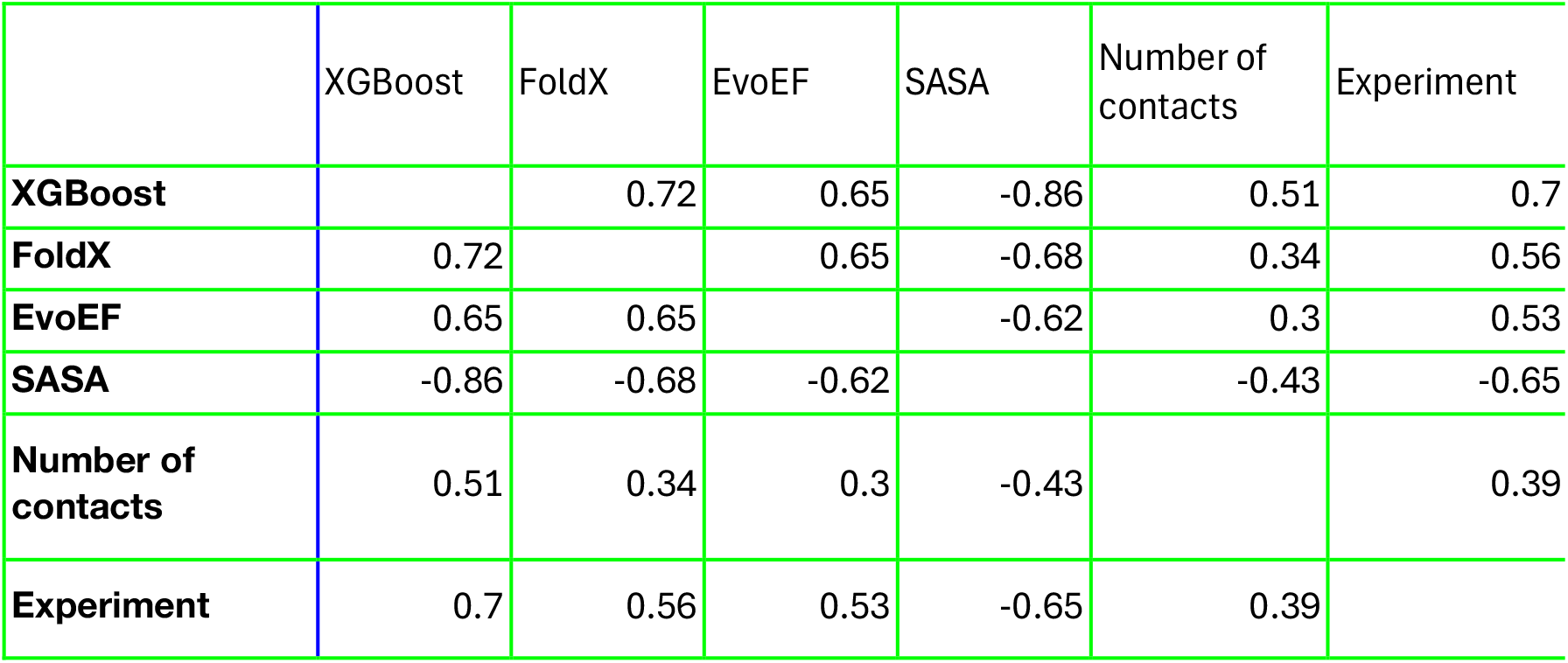
Pearson correlations for Tsuboyama et al predictions.

|  | XGBoost | FoldX | EvoEF | SASA | Number of contacts | Experiment |
| --- | --- | --- | --- | --- | --- | --- |
| **XGBoost** |  | 0.72 | 0.65 | -0.86 | 0.51 | 0.7 |
| **FoldX** | 0.72 |  | 0.65 | -0.68 | 0.34 | 0.56 |
| **EvoEF** | 0.65 | 0.65 |  | -0.62 | 0.3 | 0.53 |
| **SASA** | -0.86 | -0.68 | -0.62 |  | -0.43 | -0.65 |
| **Number of contacts** | 0.51 | 0.34 | 0.3 | -0.43 |  | 0.39 |
| **Experiment** | 0.7 | 0.56 | 0.53 | -0.65 | 0.39 |  |

**Table S2.** Pearson correlations for modeling of the pan-cancer dataset.

|  | XGBoost | phosphorylation in FoldX | FoldX (mimetic) | EvoEF | SASA |
| --- | --- | --- | --- | --- | --- |
| XGBoost |  | 0.8 | 0.88 | 0.84 | -0.87 |
| phosphorylation | 0.8 |  | 0.87 | 0.78 | -0.77 |
| FoldX | 0.88 | 0.87 |  | 0.82 | -0.82 |
| EvoEF | 0.84 | 0.77 | 0.82 |  | -0.81 |
| SASA | -0.87 | -0.77 | -0.82 | -0.81 |  |

**Table S3.**
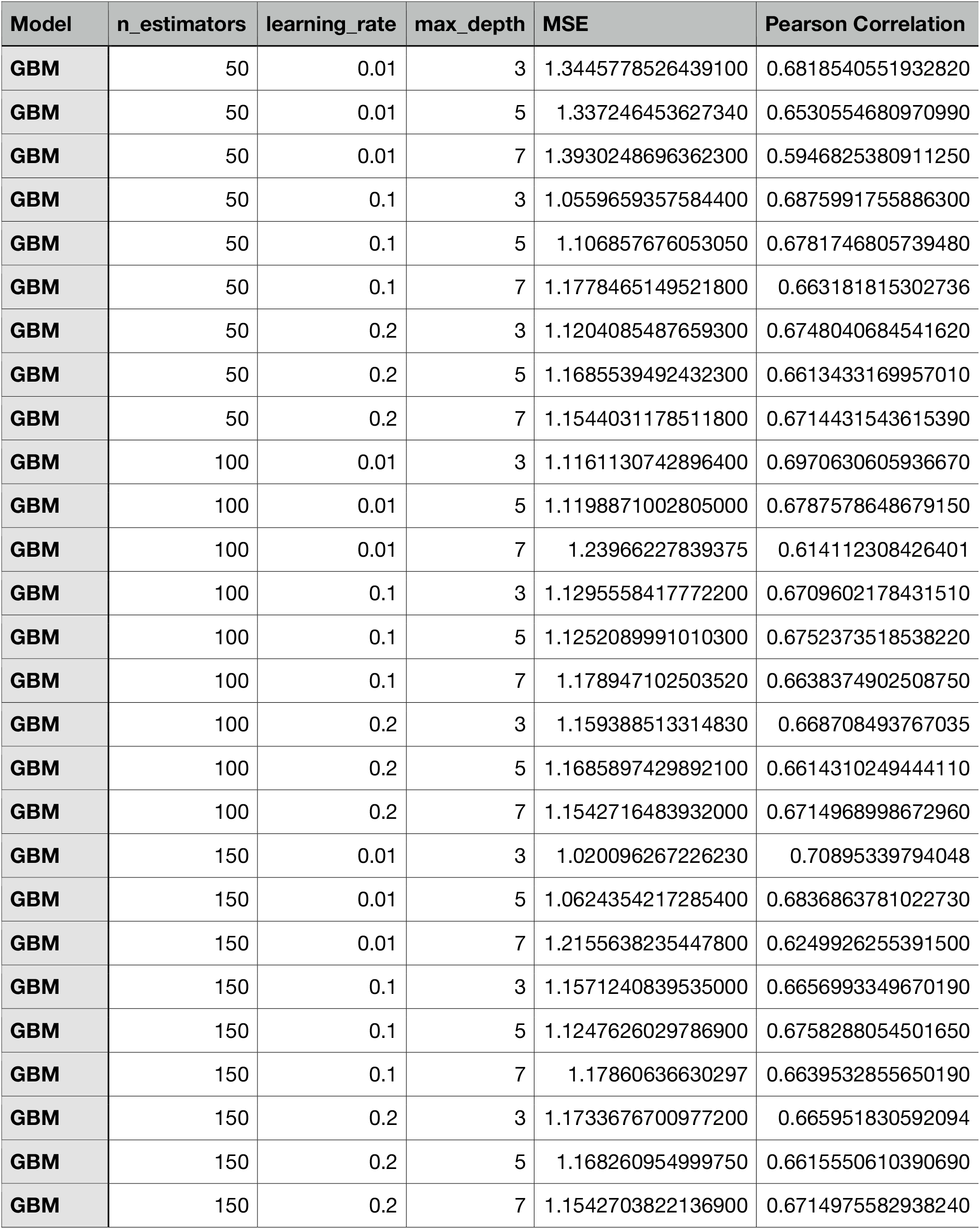
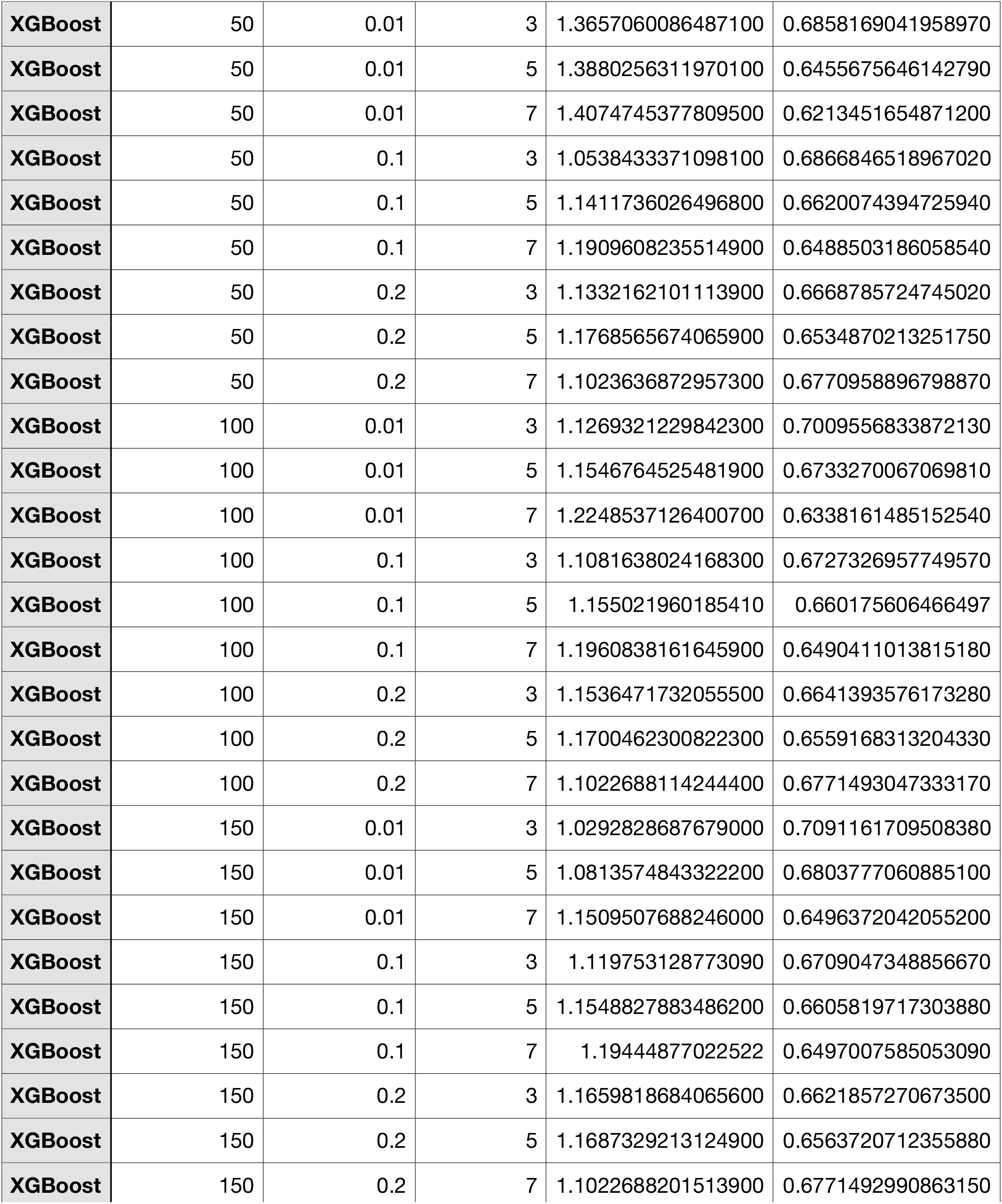
Hyperparameter optimization for model with energy calculation features.

**Table S4.**
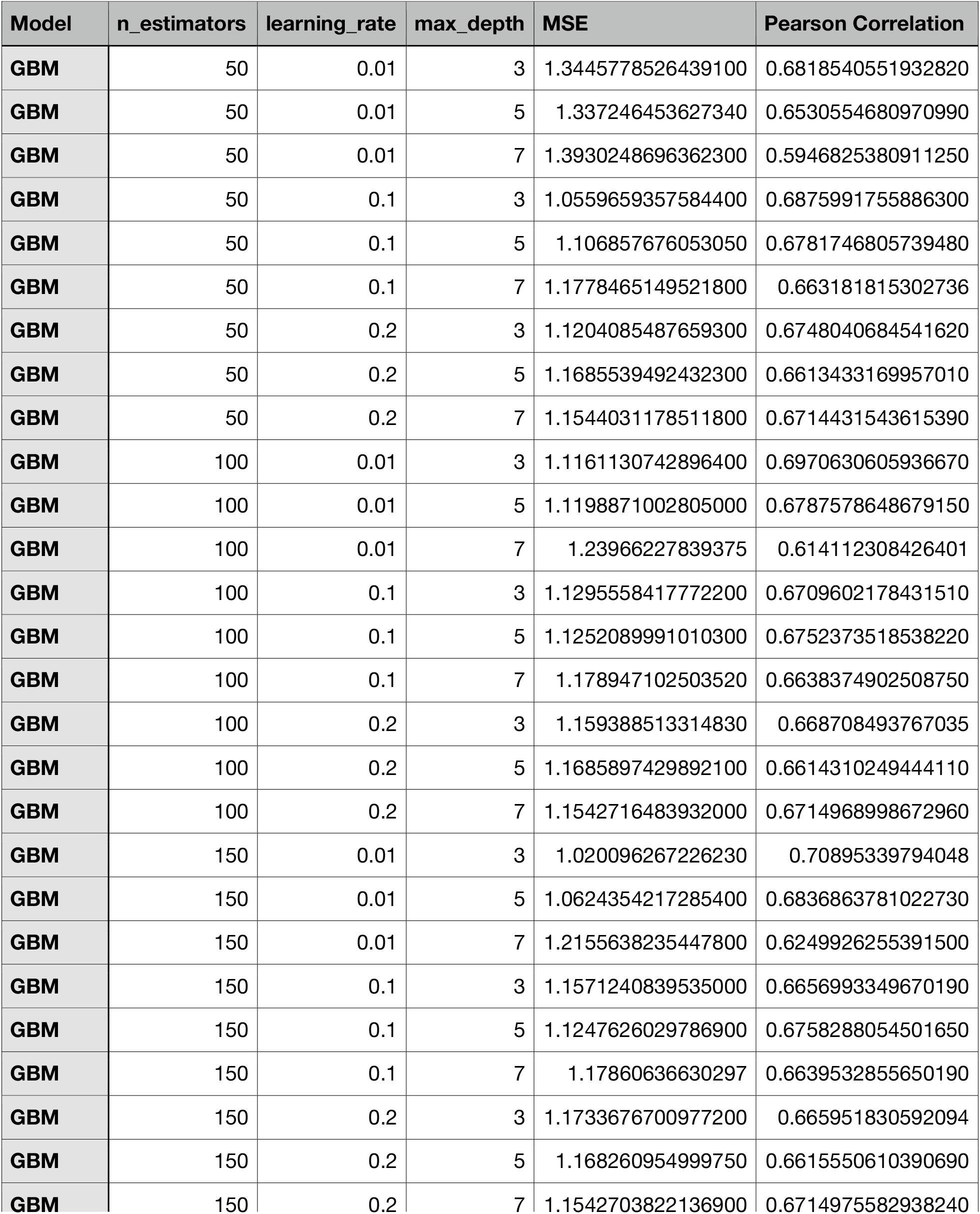
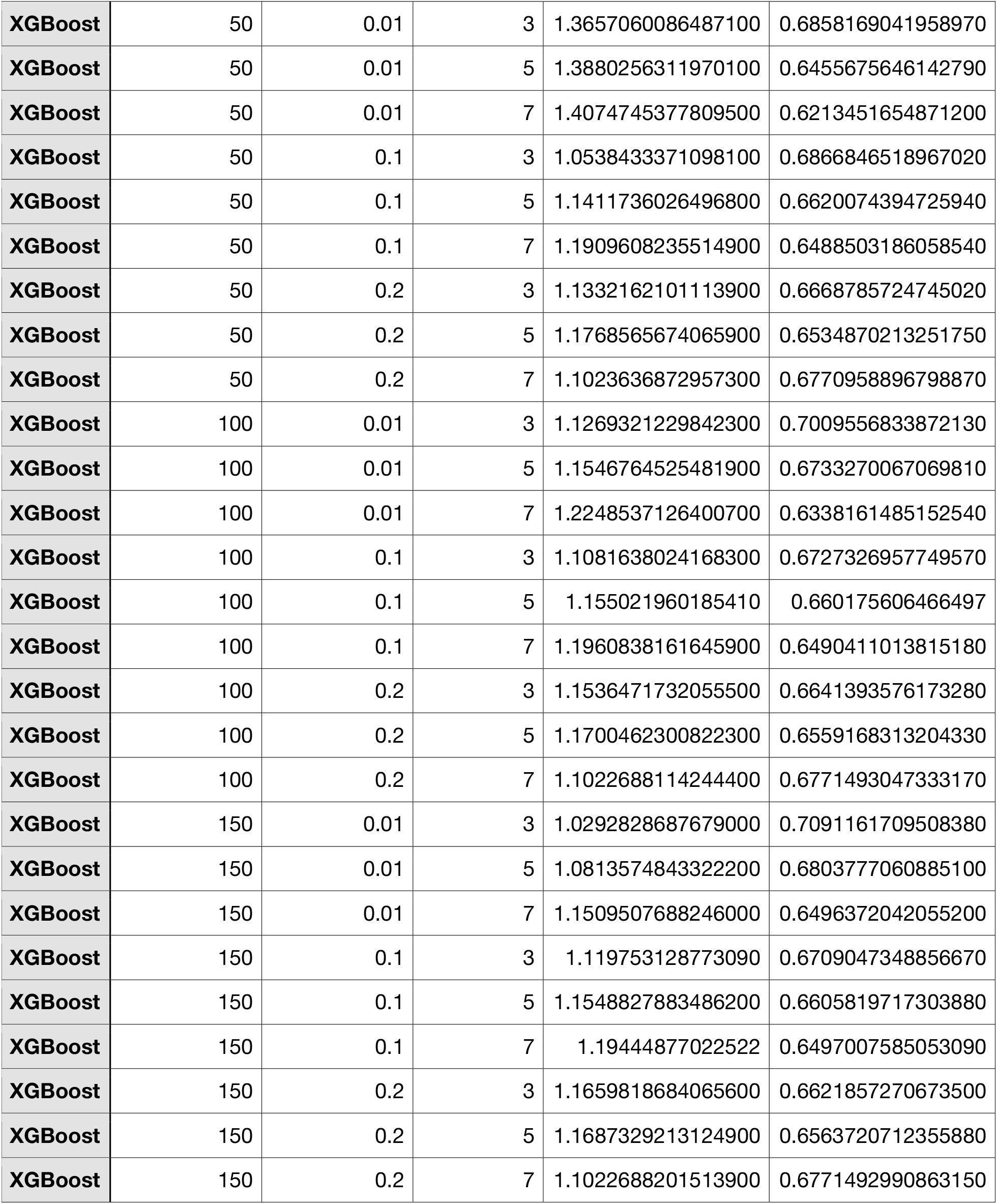
Hyperparameter optimization for rapid method without energi.

**Table S5.**
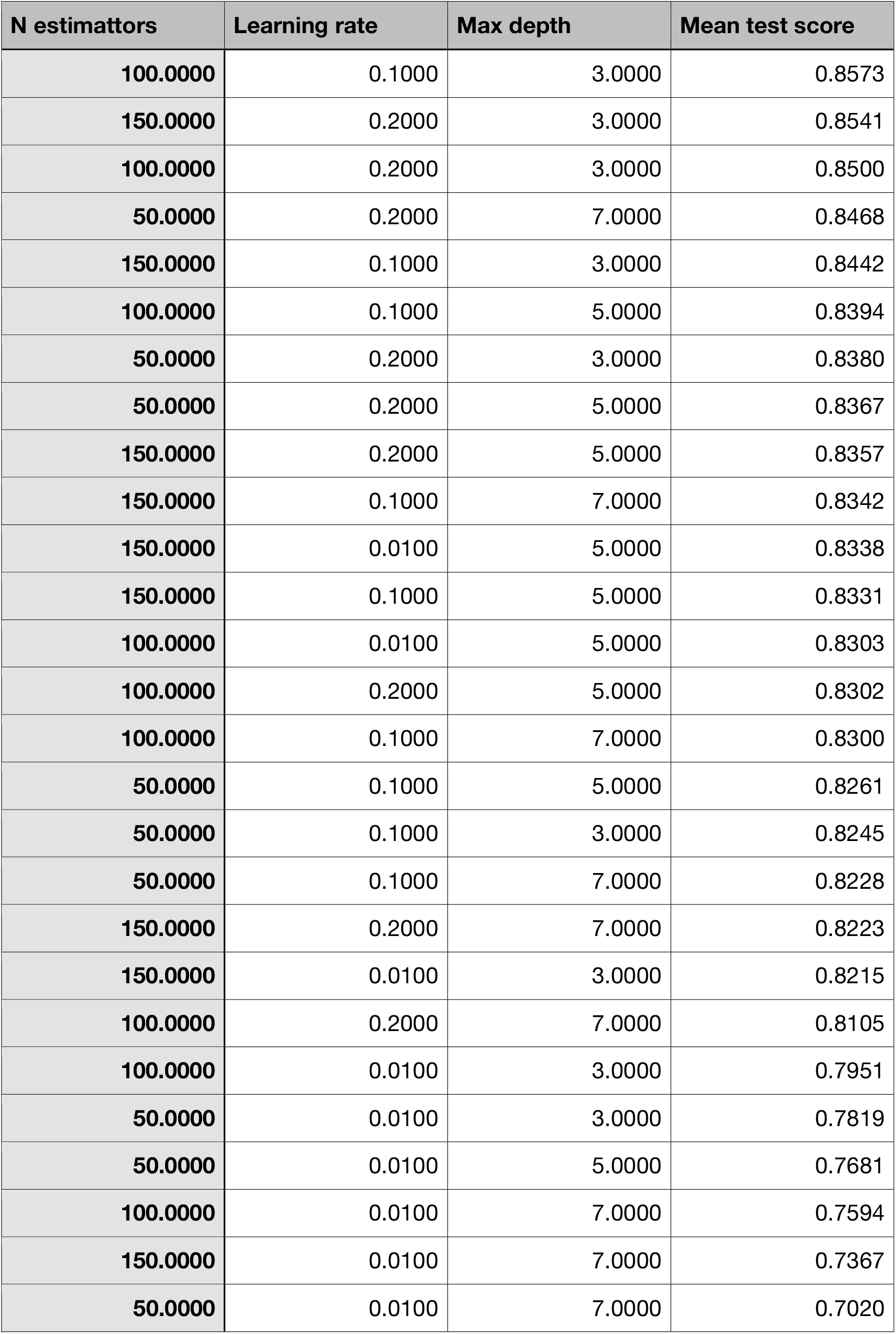
Hyperparameter optimization for the autoinhibited state vs. non-oncogene categorization.

**Table S6.** Hyperparameter optimization for oncogene vs. non-oncogene categorization.

| N estimattors | Learning rate | Max depth | Mean test score |
| --- | --- | --- | --- |
| 100.0000 | 0.2000 | 7.0000 | 0.7794 |
| 150.0000 | 0.1000 | 7.0000 | 0.7760 |
| 150.0000 | 0.2000 | 7.0000 | 0.7754 |
| 100.0000 | 0.1000 | 7.0000 | 0.7731 |
| 50.0000 | 0.1000 | 7.0000 | 0.7687 |
| 50.0000 | 0.2000 | 7.0000 | 0.7660 |
| 50.0000 | 0.2000 | 5.0000 | 0.7624 |
| 100.0000 | 0.2000 | 5.0000 | 0.7601 |
| 150.0000 | 0.2000 | 5.0000 | 0.7596 |
| 100.0000 | 0.1000 | 5.0000 | 0.7563 |
| 50.0000 | 0.1000 | 5.0000 | 0.7537 |
| 150.0000 | 0.1000 | 5.0000 | 0.7505 |
| 150.0000 | 0.0100 | 7.0000 | 0.7473 |
| 50.0000 | 0.2000 | 3.0000 | 0.7447 |
| 100.0000 | 0.2000 | 3.0000 | 0.7427 |
| 150.0000 | 0.2000 | 3.0000 | 0.7409 |
| 150.0000 | 0.1000 | 3.0000 | 0.7371 |
| 100.0000 | 0.1000 | 3.0000 | 0.7350 |
| 150.0000 | 0.0100 | 5.0000 | 0.7278 |
| 50.0000 | 0.1000 | 3.0000 | 0.7275 |
| 100.0000 | 0.0100 | 7.0000 | 0.7263 |
| 100.0000 | 0.0100 | 5.0000 | 0.7134 |
| 50.0000 | 0.0100 | 7.0000 | 0.7092 |
| 150.0000 | 0.0100 | 3.0000 | 0.6947 |
| 50.0000 | 0.0100 | 5.0000 | 0.6877 |
| 100.0000 | 0.0100 | 3.0000 | 0.6851 |
| 50.0000 | 0.0100 | 3.0000 | 0.6703 |

